# Overconfidence in visual perception in Parkinson’s disease

**DOI:** 10.1101/2020.09.09.289082

**Authors:** Orly Halperin, Roie Karni, Simon Israeli-Korn, Sharon Hassin-Baer, Adam Zaidel

**Affiliations:** Gonda Multidisciplinary Brain Research Center, Bar Ilan University, Ramat Gan 5290002, Israel; Movement Disorders Institute and the Department of Neurology, Sheba Medical Center, Tel Hashomer, Ramat Gan 5266202, Israel; The Sackler School of Medicine, Tel Aviv University, 6997801 Israel

**Keywords:** confidence, coherent motion perception, random dot kinematogram, visual, Parkinson’s disease

## Abstract

**Background:** Increased dependence on visual cues in Parkinson’s disease (PD) can unbalance the perception-action loop, impair multisensory integration, and affect everyday function of PD patients. It is currently unknown why PD patients seem to be more reliant on their visual cues.

**Objectives:** We hypothesized that PD patients may be overconfident in the reliability (precision) of their visual cues. In this study we tested coherent visual motion perception in PD, and probed subjective (self-reported) confidence in their visual motion perception.

**Methods:** 20 patients with idiopathic PD, 21 healthy aged-matched controls and 20 healthy young adult participants were presented with visual stimuli of moving dots (random dot kinematograms). They were asked to report: (1) whether the aggregate motion of dots was to the left or to the right, and (2) how confident they were that their perceptual discrimination was correct.

**Results:** Visual motion discrimination thresholds were similar (unimpaired) in PD compared to the other groups. By contrast, PD patients were significantly overconfident in their visual perceptual decisions (*p*=0.002 and *p*<0.001 vs. the age-matched and young adult groups, respectively).

**Conclusions:** These results suggest intact visual motion perception, but overestimation of visual cue reliability, in PD. Overconfidence in visual (vs. other, e.g., somatosensory) cues could underlie accounts of increased visual dependence and impaired multisensory integration in PD, and could contribute to gait and balance impairments. Future work should investigate PD confidence in somatosensory function. A better understanding of altered sensory reliance in PD might open up new avenues to treat debilitating symptoms.

## Introduction

Parkinson’s disease (PD) is a neurodegenerative disorder primarily characterized by its motor symptoms, including: bradykinesia, akinesia, muscular rigidity, resting tremor, freezing of gait (FOG) and impaired gait and balance (Almeida and Lebold, 2010; DeMaagd and Philip, 2015; Postuma *et al*., 2015). However, also non-motor symptoms manifest in PD, such as autonomic, cognitive, neurobehavioral, sleep, sensory and perceptual impairments, and these can have an important impact on a patient’s function and quality of life (Jankovic, 2008, 2017; Putcha *et al*., 2014). Among the non-motor PD symptoms, sensory and perceptual impairments may be particularly difficult to discern because they are less observable by caretakers and clinicians and can often go unnoticed even by the patient him/herself (Shulman *et al*., 2002; Chaudhuri *et al*., 2010; Bonnet *et al*., 2012).

Vision is a dominant sense in humans, and highly relied upon for everyday function. In PD, visual impairments range from basic sensory function through high-level visual processing (Weil *et al*., 2016). These include: delays in visual evoked responses (Bodis-Wollner and Yahr, 1978), contrast abnormalities (Bodis-Wollner *et al*., 1987), deterioration of spatiotemporal and color sensitivity (Montse *et al*., 2001; Weil *et al*., 2016). In addition, altered perception of visual orientation and impaired visual perception of self-motion could impact mobility, gait and balance (Davidsdottir *et al*., 2008; Gullett *et al*., 2013; Lin *et al*., 2014; Halperin *et al*., 2020; Yakubovich *et al*., 2020).

Strikingly, despite broad visual impairments, PD patients seem to demonstrate increased reliance on visual cues (Cooke *et al*., 1978; Azulay *et al*., 2002; Vaugoyeau *et al*., 2007; Davidsdottir *et al*., 2008; Barnett-Cowan *et al*., 2010; Funato *et al*., 2010). In our laboratory, we recently found that PD patients over-weighted visual self-motion cues (even though these were impaired) when integrating them with vestibular cues (Yakubovich *et al*., 2020). Thus, PD may be marked by a central brain impairment in integrating sensory information (Bertolini *et al*., 2015; Halperin *et al*., 2020; Yakubovich *et al*., 2020).

Sensory information needs to be integrated both within a specific modality and across different modalities. For example, within the visual modality, optic flow at different visual locations is integrated for coherent visual motion perception (Yuille and Grzywacz, 1988; Braddick *et al*., 2001; Kwon *et al*., 2015). Regarding the integration of cues across different modalities (multisensory integration), many recent studies have shown that cues are weighted according to their relative reliabilities (more reliable cues are more highly weighted, Jacobs, 1999; Landy and Kojima, 2001; Ernst and Banks, 2002; Alais and Burr, 2004; Fetsch *et al*., 2009; Raposo *et al*., 2012). Accordingly, the brain needs to appropriately estimate the individual cue reliabilities, while they are being sensed (Beck *et al*., 2008; Pouget *et al*., 2013). Based on reports of increased reliance on visual cues in PD, we hypothesized that PD patients might overestimate their visual cue reliability compared to their actual performance. In this study, we tested the ability to integrate visual optic flow information (coherent visual motion perception) in PD, and also probed their estimates of visual cue reliability (through subjective reports of confidence in their visual perceptual decisions).

A widely used stimulus to test coherent visual motion perception is the random dot kinematogram (RDK). RDKs present a patch of dots on a flat screen: while some of the dots move coherently in one direction (e.g., to the right or left) the remaining dots are randomly displaced. The participant’s task is to discern the overall (average) direction of motion. This becomes more difficult as the percentage of coherently moving dots is decreased. Although this is a classic and well-established paradigm that has been broadly used to test coherent visual motion perception (Morgan and Ward, 1980; Williams and Sekuler, 1984; Britten *et al*., 1992; Shadlen and Newsome, 1996; Rajananda *et al*., 2018), and has also been applied in many clinical populations, such as autism (Milne *et al*., 2002; Robertson *et al*., 2012) and Alzheimer’s disease (Fernandez and Duffy, 2012; Fernandez *et al*., 2013; Li *et al*., 2017; Song and Wang, 2019), surprisingly little is known about RDK performance in PD.

To the best of our knowledge, RDK performance in PD has been reported in only two experiments to date, neither of which found a difference in PD performance vs. controls (Putcha *et al*., 2014; Jaywant *et al*., 2016). However, Putcha *et al*. (2014) did find a positive association between increased discrimination thresholds and PD severity, and Jaywant *et al*. (2016) found an impairment in perception of biological motion from a limited amount of point lights. We also recently found impaired visual self-motion perception in PD when using an immersive virtual reality head mounted display (Yakubovich *et al*., 2020). Visual perception of biological motion and 3D visual self-motion perception (where these abovementioned PD deficits were described) require higher-level integrative brain functions. Therefore, there is still the need to test/ confirm whether or not simple (2D) coherent visual motion perception is impaired in PD. Here we tested this using RDKs.

Accordingly, in this study we address two questions: 1) is coherent visual motion perception impaired in PD? 2) Do PD patients overestimate the reliability of their visual performance? We addressed both questions using one experiment in which participants were required to discriminate the direction of visual motion (of RDKs) and then to report, on each trial their confidence in their perceptual decision. For comparison, we also tested age-matched controls as well as young adults. We found that coherent visual motion perception was unimpaired in PD (in line with previous findings; Putcha *et al*., 2014; Jaywant *et al*., 2016). However, the PD patients were significantly overconfident in their visual perceptual decisions vs. controls (despite similar performance). Biased estimates of visual cue reliability may lead to impaired multisensory integration and perceptual dysfunction in PD.

## Materials and Methods

### Participants

In this study, we tested 20 patients with idiopathic Parkinson’s disease (PD), 21 healthy aged-matched controls and 20 healthy young adult participants (participant details are presented in Suppl. Tables 1-3, respectively). The young adult group was tested in order to establish a baseline and to provide context for the PD versus age-matched control comparisons. This study was approved by the institutional Helsinki committee at The Sheba Medical Center and the internal review board at Bar-Ilan University. All participants signed informed consent before participation in the study. PD patients were recruited through the Movement Disorders Institute at Sheba Medical Center, and tested in the ‘on’ medication state, namely without any changes in their regular medication regime. Age-matched controls were recruited from the general public and staff at Bar Ilan University, and young adult participants were recruited from the student body at Bar Ilan University. Exclusion criteria included: neurological or psychiatric conditions (apart from PD), inability to walk independently or to climb stairs safely unassisted, poor corrected vision, deafness, dementia or vestibular dysfunction. Ages for the PD and age-matched control groups ranged from 32 to 80 years old. There was no significant difference in age between these two groups (*p* = 0.62; *t*-test). Ages for the young adult group ranged between 18 and 30 years. Cognitive function was assessed in all participants using the Montreal Cognitive Assessment (MoCA) test (Nasreddine *et al*., 2005).

### Stimuli and task

Participants sat 48 cm directly in front of a computer screen (DELL P2317h, 1920 × 1080 pixels). They wore headphones over their ears and held a small numeric keyboard (“numpad”) in their hands to report their answers (using the arrow keys). A schematic of the experiment is presented in Figure 1. The experiment was programmed in Python using the *Dots* (*RDK*) component (https://www.psychopy.org/builder/components/dots.html) for PsychoPy (Peirce *et al*., 2019). The visual stimuli (white moving dots) were presented within a 4.7° × 4.7° (visual angle) square aperture on a black background (6.8 dots/degree^2^ density). The visual motion stimulus for each trial lasted 1s (20 frames/s). The participants were instructed to maintain fixation during the trial on a small green fixation point, which remained static in the middle of the screen throughout the experiment. On each trial a certain percentage of the dots (% coherence) moved coherently to the right or to the left (‘signal’ dots) at 4.7°/s, while the remainder (‘noise’ dots) were randomly displaced (each noise dot was moved to a random location on every frame). Each signal dot lasted four frames (0.2s) after which it was replaced by another signal dot at a random location (at staggered/random times across dots).

**Figure 1:**
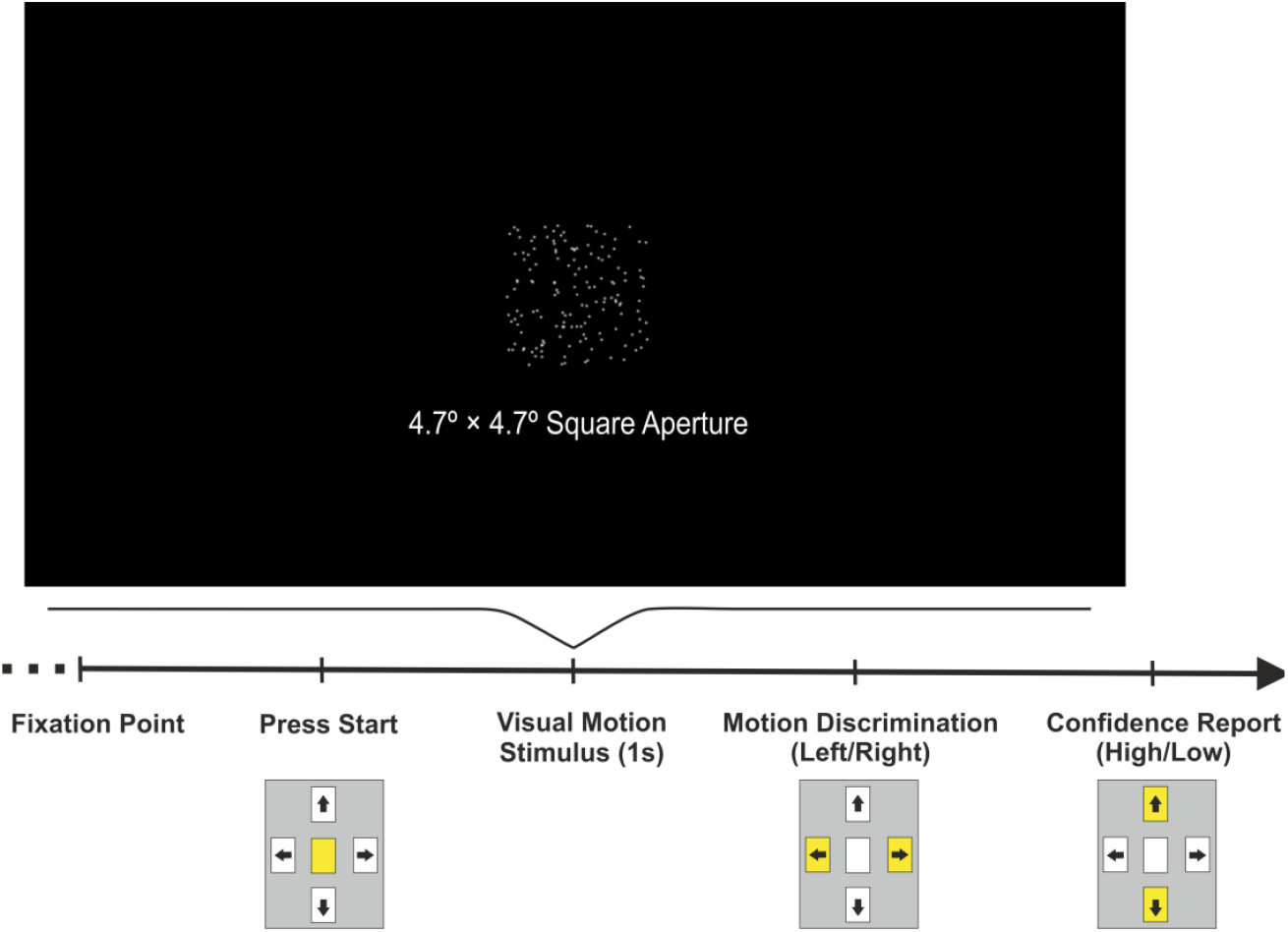
Experimental setup. The visual stimulus (white moving dots on a black screen) was presented within a 4.7° × 4.7° visual angle square aperture. The timeline (black arrow) depicts the flow of a single trial. Trials were self-initiated by pressing the center (start) button on the numpad. After the stimulus had ended, participants reported its aggregate motion direction (left button press for left and right for right) and then their level of confidence in their decision (up button press for high and down button press for low). Yellow buttons on the numpad mark possible selections at each stage.

The participants’ task was to judge whether the aggregate motion of the dots was to the right or to the left (two-alternative forced choice, 2AFC). Responses were reported by pressing the respective (right or left) arrow key on the numpad. After each visual motion discrimination, participants also reported their confidence in their choice (high or low, using the up and down arrow keys, respectively). Trials were self-paced (initiated by pressing the central button on the numpad). Different auditory tones were used to indicate i) that the system was ready for a new trial, and ii) that a choice was registered (note that the same tone was used for both incorrect and correct choices, such that no feedback was given regarding correct/incorrect choices). If a choice was not registered within a 2s window after the end of the stimulus, a response timeout was indicated by a buzzer sound (and a selection was not recorded for that trial). Participants were instructed to always make a choice (guess if unsure) and to avoid a timeout. Participants were also instructed to report their confidence right after their perceptual discrimination. The same choice tone indicated that a selection was registered, and the timeout/ buzzer was triggered if no confidence response was recorded within a 2s window since the first choice. Before starting the experiment, participants underwent a brief training period in which the experimenter gave verbal feedback in order to confirm that they understood the instructions and performed the task adequately.

On each trial the direction of motion was selected randomly (*p =* 0.5 to the right/left). We mark leftward directions by a negative sign and rightward motions by a positive sign (e.g., −100% marks 100% coherent motion to the left). Stimulus difficulty began at ±100% coherence (the easiest level with all dots moving in the same direction) and adapted individually to each participant’s performance according to a staircase procedure, as follows: if the participant gave a correct answer on the previous trial, the coherence magnitude (absolute value) was decreased by a factor of 2 (i.e., became harder) with *p =* 0.3 (and remained at the same level with *p =* 0.7). If the participant gave an incorrect answer on the previous trial, the coherence magnitude was increased by a factor of 2 (i.e., became easier) with *p* = 0.8 (and remained at the same level with *p* = 0.2). This staircase rule converges at ∼73% correct responses (i.e., the same level of success/ subjective difficulty across participants). A major advantage of this method here (when testing confidence) is that the success rate (accuracy) is similar for all participants (Fleming *et al*., 2010). Thus, any differences in confidence do not reflect differences in success. Each participant performed 150 trials (which took ∼13 minutes).

### Data analyses and statistics

The data analyses were performed with custom software using Matlab R2013b (The MathWorks) and the psignifit toolbox for Matlab version 4 (Schütt *et al*., 2016). Two psychometric plots were constructed for each participant. (i) The first plot depicts each participant’s psychophysics performance in visual motion discrimination (see examples in Figure 2A). This was quantified by the rightward choice ratio as a function of coherence. The data were fit (per participant) with a cumulative Gaussian distribution function, and the psychophysical ‘threshold’ defined by the standard deviation (SD) of the fitted curve. Better performance (i.e., higher reliability) is reflected by a lower threshold. (ii) The second plot represents the confidence reports (see examples in Figure 3A). This was quantified by the ratio of high-confidence reports as a function of coherence. The confidence reports for leftward motion choices were flipped (to the symmetrical coherence) so that the data could be pooled and depicted by one plot. We fit these data (circle markers) with a cumulative Gaussian distribution function, and defined the confidence bias by the value at which the curve crosses 0% coherence (i.e., the case for which there was no objective motion information in the signal). Statistical analyses were performed using JASP (version 0.10.2) and Matlab (details of which are presented together with the relevant results).

**Figure 2:**
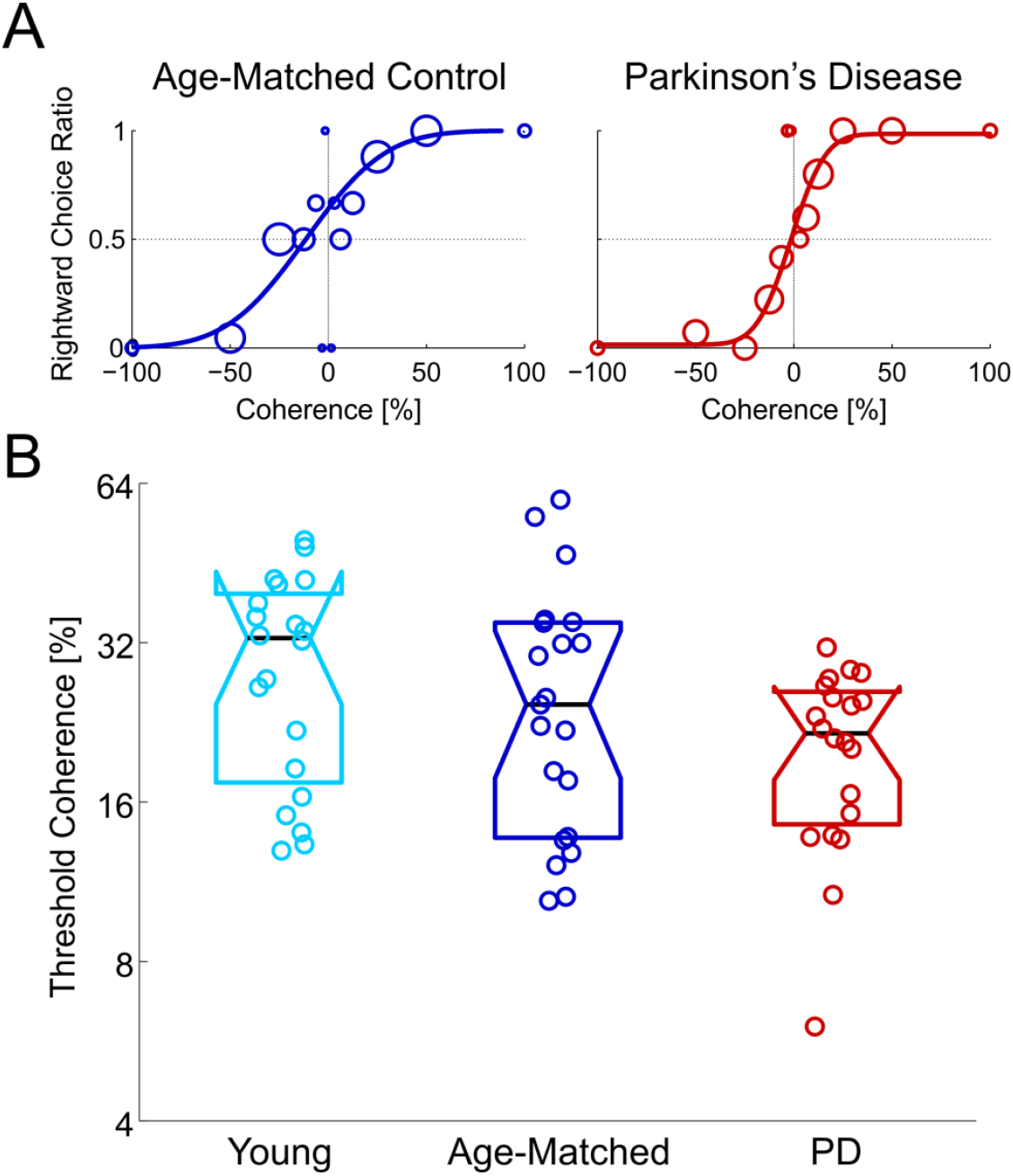
Visual motion perception thresholds are unimpaired in PD. (A) Visual motion discrimination psychometric plots for an example age-matched control participant (dark blue) and an example PD patient (red). Circle markers represent the ratio of rightward choices per coherence. The sizes of the circle markers reflect the number of trials for each data point. The data were fitted with cumulative Gaussian distribution functions (solid lines). (B) Visual motion perception thresholds for the young adult (light blue), age-matched (dark blue) and PD (red) groups. Each data point (jittered slightly horizontally for visibility) marks the threshold of a single participant. The black horizontal line in each boxplot indicates the group median. Overlapping notches between the boxes indicate that the medians do not differ at the *p* = 0.05 significance level.

**Figure 3:**
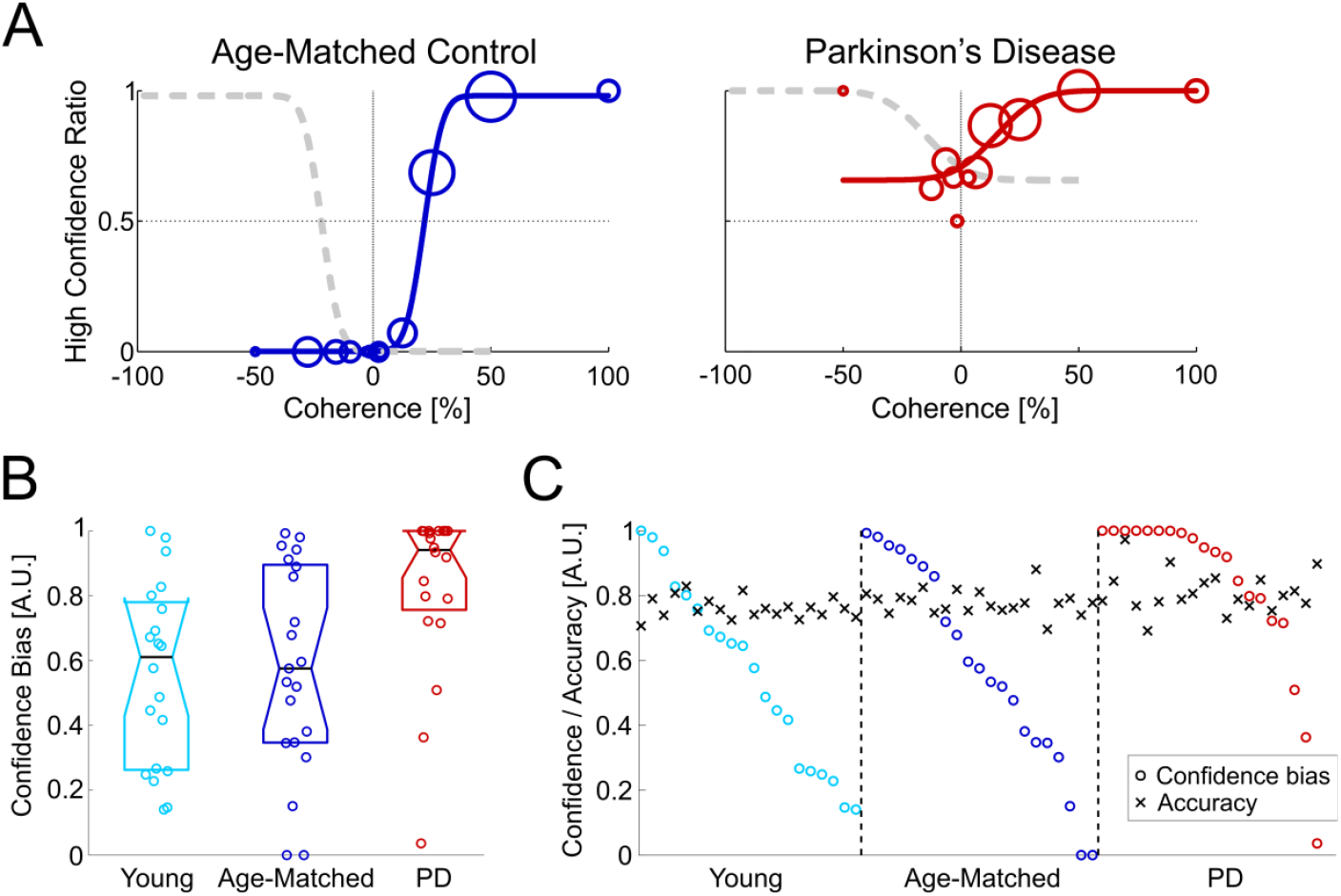
Visual overconfidence in PD. (A) Confidence plots for an example age-matched control participant (dark blue) and an example PD patient (red). Circle markers represent the ratio of high confidence selections for rightward choices (data for leftward choices were flipped to the symmetrical coherences for pooling). The sizes of the circle markers reflect the number of trials for each data point. The data were fitted with cumulative Gaussian distribution functions (solid lines). The light gray dashed lines reflect the mirror image of the Gaussian fits (to depict the same plot when oriented to leftward choices). (B) Confidence biases are presented for the young adult (light blue), age-matched (dark blue) and PD (red) groups. Each data point (jittered slightly horizontally for visibility) marks the high-confidence ratio bias at 0% coherence, per participant. The black horizontal line in each boxplot indicates the group median. Non-overlapping notches of the PD vs. the other two groups indicates that the PD median differs from the other two at the *p* = 0.05 significance level (notches for the two non-PD groups overlap). (C) The high-confidence ratio bias at 0% coherence (sorted per group by descending value) and corresponding performance accuracy (correct choice ratio; black ‘x’ markers), per participant.

## Results

### Intact 2D coherent visual motion perception in PD

Psychometric curves of visual motion discrimination are presented for an example PD patient and an example age-matched control participant in Figure 2A (red and blue curves, respectively). A steeper psychometric curve marks better performance (higher cue reliability). This was quantified by the ‘threshold’ (SD of the fitted cumulative Gaussian function), where lower values reflect better performance. In these examples, the PD participant had a better visual motion perception threshold (14% coherence) vs. the control participant (32% coherence).

At the group level (Fig. 2B), visual motion coherence thresholds in PD were comparable to the other groups. There was no significant difference across groups (*p =* 0.073, ANOVA; a Bayesian ANOVA did not provide evidence for, or against, the null hypothesis, BF_10_ = 0.98). If anything, PD thresholds were marginally better (lower). Therefore these results indicate that PD patients have intact coherent visual motion perception. We also did not find any significant correlations between thresholds and clinical measures of disease severity or progression: UPDRS (‘on’ or ‘off’ medication), levodopa equivalent daily dose (LEDD; Tomlinson *et al*., 2010), or disease duration (*p* > 0.05 for all, before correction for multiple comparisons). These results suggest that PD deficits in complex visual motion paradigms, including biological motion (Jaywant *et al*., 2016) and 3D visual self-motion perception (Yakubovich *et al*., 2020) likely reflect more complex impairments of higher-level brain functions for interpreting visual motion, and not a basic visual motion perception deficit in PD.

### Visual overconfidence in PD

Participants were required (after each visual discrimination) to report their confidence (high or low) regarding their visual discrimination response. The ratio of high-confidence responses was calculated per coherence. Confidence curves for the same two example participants in Figure 2A are presented in Figure 3A. These confidence data are presented in terms of rightward choices, but also comprise confidence reports for leftward choices. Specifically, the polarity of the stimulus (coherence sign) was flipped for leftward choices, in order to pool with rightward choices. For example, confidence reports for leftward choices at - 100% coherence were pooled with confidence reports for rightward choices at 100% coherence (both easy stimuli and correct choices). Similarly, confidence reports for incorrect leftward choices (e.g., at 12.5% coherence) were pooled with those for symmetrical (incorrect) rightward choices (e.g., at −12.5% coherence). Accordingly, in these pooled confidence curves negative coherences represent incorrect choices (a rightward choice for a negative stimulus or a leftward choice for a positive stimulus) and positive coherences represent correct choices.

For the example control participant (Fig 3A, left) confidence is high (close to 1) for easy stimuli (50-100% coherence), and low (close to 0) for incorrect choices (negative coherence values). By contrast, although the PD participant (Fig 3A, right) also reports high confidence for the easy stimuli (50-100% coherence), confidence only drops off slightly for negative coherences (incorrect choices), and still remains high. The y-intercept (where the curve crosses 0% coherence) reflects the high-confidence ratio for an ambiguous stimulus. We used this value as a measure of the confidence bias. For the example control participant, the y-intercept is close to 0. This indicates low confidence in visual discriminations around the ambiguous stimuli. By contrast, the y-intercept for the PD patient is 0.72 – indicating a high-confidence bias for ambiguous stimuli.

There was a significant difference in confidence bias (y-intercept of the fitted curves) across the three groups (Fig. 3B; *p* = 0.002; Kruskall-Wallis). Post-hoc comparisons (Dunn’s) revealed significantly higher values in PD vs. both age-matched controls (*p* = 0.002) and young adults (*p* = 6.1·10^−4^), and no significant difference between the two non-PD groups (*p* = 0.38, BF_10_ = 0.31). These results indicate that PD patients are overconfident regarding their performance in coherent visual motion perception. We did not find any significant correlations between this bias in confidence and clinical measures: UPDRS (‘on’ or ‘off’ medication), levodopa equivalent daily dose (LEDD; Tomlinson *et al*., 2010), or disease duration (*p* > 0.05 for all, before correction for multiple comparisons).

There were six participants (all PD) whose confidence bias was equal to 1 (meaning that they always chose high-confidence). During training, when we noticed that a participant was only choosing one confidence option (e.g., high confidence), we stopped to explain the task again, and provided further training, during which we insisted that they also choose the other option (e.g., low confidence) in order to demonstrate that they understood our instructions. Only when participants did this, and we confirmed that they fully understood the task, did we proceed with the actual experiment (these participants also confirmed verbally during this interaction that they were indeed highly confident in all their choices). During the actual experiment we did not interfere or comment further on their confidence choices so as not to bias the results. Due to these training measures (and the observation that confidence bias values of exactly one were seen only in the PD group) we interpret these results as a real confidence bias in PD (and thus did not exclude these participants from the analysis). Nonetheless, even when removing these six PD participants, the Dunn’s comparisons remain significant (albeit more marginally) between PD vs. age-matched controls (*p* = 0.04) and PD vs. young adults (*p* = 0.02).

### Confidence bias for comparable accuracies

In our experiment, we used a staircase procedure that adapts task difficulty according to each individual’s correct/incorrect responses. This procedure maintains similar accuracy (proportion of correct responses) across participants. Note that accuracy is not the same as reliability, which (as described above) relates to the steepness of the psychometric curve (Zaidel *et al*., 2011). Hence, individuals can have different cue reliability (measured by thresholds, Fig. 2B) but similar accuracy (controlled by the staircase). Since accuracy was maintained similar across participants (∼73%), differences in confidence do not result from different levels of success/accuracy.

Figure 3C presents the confidence bias (colored circles) for each participant sorted by descending value (per group) together with their respective accuracy (black ‘x’ markers). The confidence bias values are largely heterogeneous across individuals (but with more high-confidence values for PD), even though accuracy was largely consistent. To further test this point, we filtered out participants whose accuracy was > 80% (this could occur if a participant didn’t respond to some difficult trials, which were therefore not included in the analysis). Also when analyzing only those participants whose accuracy was lower than 80% (Fig. S1) confidence values were still significantly different across the three groups (*p* = 0.006, Kruskall-Wallis) and significantly higher in PD vs. both age-matched controls (*p* = 0.003; Dunn’s Post-hoc comparison) and young adults (*p* = 0.001; Dunn’s Post-hoc comparisons).

### Cognitive function

MoCA scores differed significantly across the three groups (*p* = 3.3·10^−6^, ANOVA) with PD having lower scores (mean ± SD = 23.2 ± 3.0) vs. both age-matched controls (26.3 ± 1.7; *p* = 2.8·10^−4^, Tukey posthoc) and young adults (27.3 ± 2.3; *p* = 4.5·10^−6^, Tukey posthoc). This does not likely account for any of our findings because MoCA scores did not correlate significantly with confidence bias (*R* = −0.16, *p* = 0.21, BF_10_ = 0.35) or thresholds (*R* = 0.15, *p* = 0.26, BF_10_ = 0.30). Furthermore, an ANCOVA with MoCA as a covariate still found a significant difference in confidence bias scores between groups (*p* = 0.012) with PD patients remaining significantly overconfident vs. age-matched controls (*p* = 0.021, Tukey posthoc) and young adults (*p* = 0.018, Tukey posthoc).

## Discussion

In this study we found unimpaired coherent visual motion perception in PD. This in line with two other recent studies (Putcha *et al*., 2014; Jaywant *et al*., 2016). However, despite similar psychophysical performance, PD patients were significantly overconfident in their perceptual decisions. This suggests that PD patients overestimate the reliability of their visual cues, which might explain recent findings of visual overweighting during multisensory integration (Yakubovich *et al*., 2020), and descriptions of increased dependence on vision (Cooke *et al*., 1978; Bronstein *et al*., 1990; Azulay *et al*., 1999, 2002; Almeida and Lebold, 2010; Cowie *et al*., 2010) in PD.

White et al. (2016) recently found that PD patients overestimate their olfactory performance (in spite of olfactory impairment). Accordingly, PD overconfidence in perceptual function is not limited to one sensory domain. By contrast, self-reported confidence in mobility in PD predicts the risk of falling (Mak and Pang, 2009) and is correlated with specific characteristics of gait and balance function (Curtze *et al*., 2016). Hence, PD patients do seem to be aware of impaired balance control. Together, these results suggest that the metacognitive ability to assess perceptual function may differ across sensory modalities in PD. Since effective multisensory integration requires good estimates of the individual cue reliabilities relative to one-another, this could lead to impaired multisensory integration in PD, depending on which cues are being integrated. Appropriate estimates of balance function (Mak and Pang, 2009) together with overconfidence in visual function (described here) might together explain visual overweighting in PD during vestibular-visual integration (Yakubovich *et al*., 2020). Further research that specifically tests PD confidence in vestibular performance (in relation to actual vestibular function) compared to vision is needed to confirm this.

In terms of visual motion perception, our findings here (and those of Putcha *et al*., 2014 and Jaywant *et al*., 2016) suggest that basic (2D) visual motion perception is unimpaired in PD. By contrast, we recently found increased thresholds of visual self-motion perception in PD when using an immersive virtual reality head mounted display (Yakubovich *et al*., 2020). These self-motion stimuli differ from RDKs in that they are rendered in 3D, cover a wide field-of-view, follow a Gaussian motion profile (soft start and end) and are presented in the context of vestibular (inertial) motion – vs. RDKs, which span a narrow field-of-view, are presented in 2D (on a flat screen), with constant motion speed, and thus specifically test perception of visual motion (rather than self-motion). These results, together with Jaywant *et al*. (2016) who found impaired biological motion perception in PD, suggest that impairments of higher-level visual functions do not result from low-level visual impairments (since basic visual motion perception seems intact). Rather these may reflect integration deficits in higher order cortical areas (Halperin *et al*., 2020).

Our results here (and those of Putcha *et al*., 2014 and Jaywant *et al*., 2016) were collected in the ‘on’ medication state. Future studies should investigate coherent visual motion perception also in the ‘off’ medication state. Curtze et al. (2016) found that confidence reports in PD better predict objective measures of mobility in the ‘off’ (vs. ‘on’) medication state. This may suggest that metacognitive mechanisms to estimate ones function in the ‘on’ state are inaccurate. Hence, perceptual confidence in PD requires further investigation in future studies that compare perceptual confidence (vs. function) both ‘on’ and ‘off’ medication.

We found here that for comparable visual motion perceptual function, self-estimates of this function were larger in PD. This suggests that PD patients overestimate the reliability of their visual cues (this might be specific to vision or general). This can be dangerous, especially in situations of impaired function (such as higher-level integrative visual function). Furthermore, it can lead to over-reliance on visual cues during multisensory integration, which would further impair function. Over-reliance on vision vs. somatosensory (e.g. vestibular and perceptual) cues can affect gait and balance, which are debilitating and difficult to treat symptoms in PD. A better understanding of these mechanisms might open up new avenues for alternative therapies to better treat these symptoms, such as sensorimotor retraining techniques.

## Acknowledgements

We would like to thank Avraham Elkaras for programming assistance, Tamar Harpaz for management assistance, and the staff of the Movement Disorders Institute at Sheba Medical Center for help with patient recruitment. We would also like to thank Roy Salomon for constructive advice in preparing the paper.

## Authors’ roles

Adam Zaidel conceived of and oversaw the research project. Orly Halperin performed the experiments. Orly Halperin and Roie Karni performed the analyses. Orly Halperin and Adam Zaidel wrote the manuscript. Simon Israeli-Korn and Sharon Hassin-Baer cared for and recruited the Parkinson’s patients and reviewed and critiqued the manuscript.

## Supplementary figure

**Figure S1.**
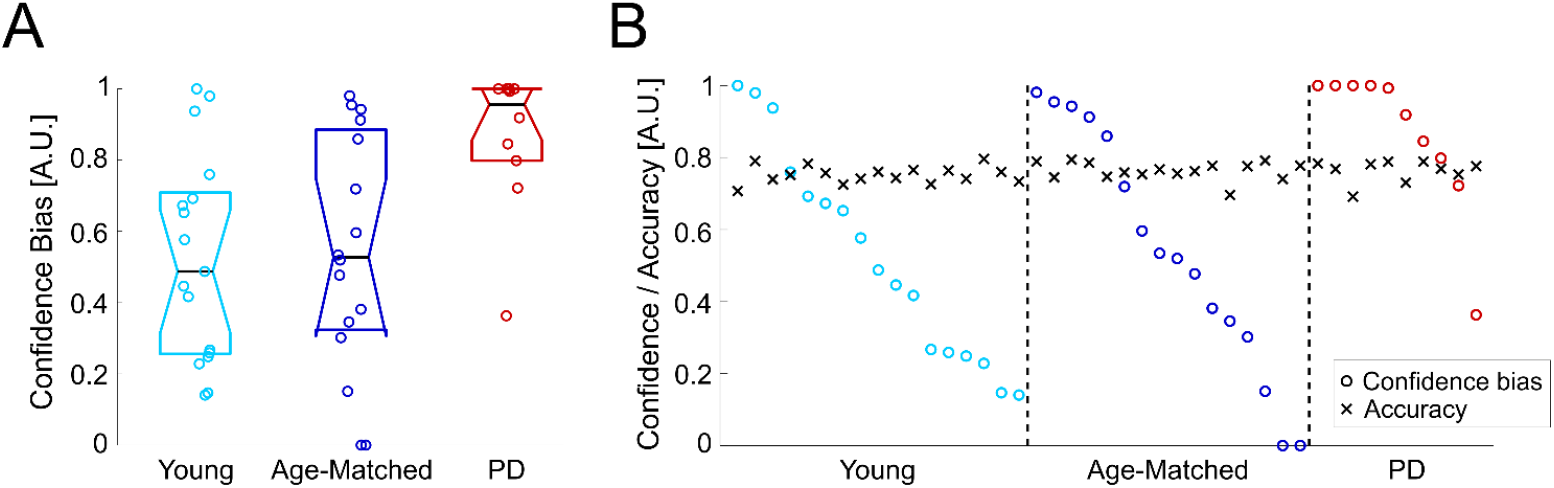
Visual overconfidence in PD also after removing participants with accuracy > 0.8. All conventions are the same as Figure 3B and C. Results remain the same (overconfidence in PD) even after removing participants with accuracy > 0.8 (N = 17, 16 and 10 for the young, age-matched and PD groups remained, respectively).

**Supplementary Table 1:** Parkinson’s disease participants’ details

| Subject ID | Age (years) | Gender | MoCA | First symptom | Side of first symptom | Disease duration (years) | Levodopa initiation (year) | Motor UPDRS ON | Motor UPDRS OFF | LEDD (mg) |
| --- | --- | --- | --- | --- | --- | --- | --- | --- | --- | --- |
| PD1 | 76 | F | 18 | Toe tremor | Right | 7 | 2015 | 16 | 15 | 400 |
| PD2 | 70 | F | 23 | Hand slowness, handwriting less clear | Right | 7 | LN | 34 | 45 | 300 |
| PD3 | 65 | M | 23 | Hand tremor | Left | 4 | LN | 20 | - | 300 |
| PD4 | 58 | M | 23 | Hand tremor | Right | 17 | 2007 | 34 | 44 | 3120 |
| PD5 | 79 | M | 25 | Leg tremor | Right | 11 | 2016 | 17 | 18 | 550 |
| PD6 | 75 | M | 22 | Leg tremor | Right | 13 | 2017 | 10 | 17 | 487.5 |
| PD7 | 68 | M | 17 | Hand tremor | Left | 13 | 2007 | 31 | 51 | 750 |
| PD8 | 64 | F | 28 | Hand tremor | Left | 2 | LN | 11 | - | 100 |
| PD9 | 32 | F | 29 | Leg dragging | Right | 3 | LN | 17 | - | 100 |
| PD10 | 52 | M | 23 | Hand dysfunction | Left | 3 | LN | 35 | - | 350 |
| PD11 | 44 | F | 24 | Hand dysfunction | Right | 16 | 2011 | 15 | 39 | 815 |
| PD12 | 70 | M | 21 | Hand tremor | Left | 11 | 2009 | 25 | 46 | 1298.5 |
| PD13 | 65 | M | 19 | Reduced arm swing | Right | 6 | 2014 | 21 | - | 350 |
| PD14 | 62 | M | 25 | Hand dysfunction | Left | 14 | 2010 | 22 | 29 | 532 |
| PD15 | 66 | M | 22 | Reduced arm swing | Right | 2 | LN | 32 | - | 100 |
| PD16 | 70 | M | 22 | Mouth tremor | Left | 3 | 2017 | 23 | 24 | 650 |
| PD17 | 54 | M | 27 | Hand dysfunction | Left | 9 | 2014 | 31 | 24 | 499 |
| PD18 | 82 | M | 25 | Hand tremor | Left | 6 | 2017 | 31 | - | 325 |
| PD19 | 72 | F | 25 | Hand tremor | Right | 6 | LN | 33 | - | 350 |
| PD20 | 80 | M | 23 | Hand tremor | Right | 1 | 2019 | 41 | - | 300 |
| **Mean ± SD** | **65.2 ± 12.1** | **14M:6F** | **23.2 ± 3.0** |  |  | **7.7 ± 5.0** |  | **25.0 ± 9.0** | **32.0 ± 13.3** | **583.9 ± 659.2** |

MoCA: Montreal cognitive assessment (range 0 to 30, higher scores reflect better cognition); UPDRS: Unified Parkinson's Disease Rating Scale (motor part, range 0 to 108, higher scores reflects more severe motor symptoms, ON = when taking antiparkinsonian medication, OFF = when not taking antiparkinsonian medication); PD: Parkinson's disease participant; LN: levodopa naïve; LEDD: levodopa equivalent daily dose of medication (according to Tomlinson *et al.*, 2010).

**Supplementary Table 2:** Age-matched control participants’ details

| **Subject ID** | **Age (years)** | **Gender** | **MoCA** |
| --- | --- | --- | --- |
| AMC1 | 64 | F | 28 |
| AMC2 | 66 | M | 27 |
| AMC3 | 63 | F | 28 |
| AMC4 | 65 | F | 28 |
| AMC5 | 54 | M | 27 |
| AMC6 | 68 | F | 28 |
| AMC7 | 52 | F | 27 |
| AMC8 | 69 | M | 25 |
| AMC9 | 69 | M | 25 |
| AMC10 | 67 | M | 26 |
| AMC11 | 54 | F | 27 |
| AMC12 | 65 | F | 25 |
| AMC13 | 62 | F | 25 |
| AMC14 | 62 | F | 26 |
| AMC15 | 63 | F | 25 |
| AMC16 | 71 | M | 26 |
| AMC17 | 64 | F | 25 |
| AMC18 | 61 | F | 30 |
| AMC19 | 73 | F | 23 |
| AMC20 | 71 | F | 24 |
| AMC21 | 54 | M | 28 |
| **Mean ± SD** | **63.7 ± 6.0** | **7M:14F** | **26.3 ± 1.7** |

MoCA: Montreal cognitive assessment (range 0 to 30, higher scores reflect better cognition); AMC: age-matched control participant.

**Supplementary Table 3:** Young adult participants’ details

| **Subject ID** | **Age (years)** | **Gender** | **MoCA** |
| --- | --- | --- | --- |
| YA1 | 30 | F | 25 |
| YA2 | 29 | F | 25 |
| YA3 | 28 | M | 30 |
| YA4 | 25 | F | 27 |
| YA5 | 18 | F | 28 |
| YA6 | 23 | M | 26 |
| YA7 | 22 | F | 29 |
| YA8 | 28 | M | 23 |
| YA9 | 29 | M | 27 |
| YA10 | 29 | M | 27 |
| YA11 | 21 | F | 27 |
| YA12 | 25 | M | 30 |
| YA13 | 27 | M | 27 |
| YA14 | 23 | F | 30 |
| YA15 | 25 | F | 29 |
| YA16 | 23 | M | 30 |
| YA17 | 26 | M | 25 |
| YA18 | 24 | M | 23 |
| YA19 | 22 | F | 27 |
| YA20 | 26 | F | 30 |
| **Mean ± SD** | **25.2 ± 3.2** | **10M:10F** | **27.3 ± 2.3** |

MoCA: Montreal cognitive assessment (range 0 to 30, higher scores reflect better cognition); YA: young adult participant.

## References

Alais D, Burr D. The ventriloquist effect results from near-optimal bimodal integration. Curr. Biol 2004; 14: 257–262.

Almeida QJ, Lebold CA. Freezing of gait in Parkinson’s disease: a perceptual cause for a motor impairment? J. Neurol. Neurosurg. Psychiatry 2010; 81: 513–518.

Azulay JP, Mesure S, Amblard B, Blin O, Sangla I, Pouget J. Visual control of locomotion in Parkinson’s disease. Brain 1999; 122: 111–120.

Azulay JP, Mesure S, Amblard B, Pouget J. Increased visual dependence in Parkinson’s disease. Percept. Mot. Ski. 2002; 95: 1106–1114.

Barnett-Cowan M, Dyde RT, Fox SH, Moro E, Hutchison WD, Harris LR. Multisensory determinants of orientation perception in Parkinson’s disease. Neuroscience 2010; 167: 1138–1150.

Beck JM, Ma WJ, Kiani R, Hanks T, Churchland AK, Roitman J, et al. Probabilistic population codes for Bayesian decision making. Neuron 2008; 60: 1142–1152.

Bertolini G, Wicki A, Baumann CR, Straumann D, Palla A. Impaired tilt perception in Parkinson’s disease: a central vestibular integration failure. PLoS One 2015; 10: e0124253.

Bodis-Wollner I, Marx MS, Mitra S, Bobak P, Mylin L, Yahr M. Visual dysfunction in parkinson’s disease: Loss in spatiotemporal contrast sensitivity. Brain 1987; 110: 1675–1698.

Bodis-Wollner I, Yahr MD. Measurements of visual evoked potentials in Parkinson’s disease. Brain a J. Neurol. 1978; 101: 661–671.

Bonnet AM, Jutras MF, Czernecki V, Corvol JC, Vidailhet M. Nonmotor symptoms in Parkinsons disease in 2012: Relevant clinical aspects. Parkinsons. Dis. 2012

Braddick OJ, O’Brien JMD, Wattam-Bell J, Atkinson J, Hartley T, Turner R. Brain areas sensitive to coherent visual motion. Perception 2001; 30: 61–72.

Britten KH, Shadlen MN, Newsome WT, Movshon JA. The analysis of visual motion: A comparison of neuronal and psychophysical performance. J. Neurosci. 1992; 12: 4745–4765.

Bronstein AM, Hood JD, Gresty MA, Panagi C. Visual control of balance in cerebellar and parkinsonian syndromes. Brain 1990; 113: 767–779.

Chaudhuri KR, Prieto-Jurcynska C, Naidu Y, Mitra T, Frades-Payo B, Tluk S, et al. The nondeclaration of nonmotor symptoms of Parkinson’s disease to health care professionals: an international study using the nonmotor symptoms questionnaire. Mov. Disord. 2010; 25: 704–709.

Cooke JD, Brown JD, Brooks VB. Increased Dependence on Visual Information for Movement Control in Patients with Parkinson’s disease. Can. J. Neurol. Sci. / J. Can. des Sci. Neurol. 1978; 5: 413–415.

Cowie D, Limousin P, Peters A, Day BL. Insights into the neural control of locomotion from walking through doorways in Parkinson’s disease. Neuropsychologia 2010; 48: 2750–2757.

Curtze C, Nutt JG, Carlson-Kuhta P, Mancini M, Horak FB. Objective Gait and Balance Impairments Relate to Balance Confidence and Perceived Mobility in People With Parkinson Disease. Phys. Ther. 2016

Davidsdottir S, Wagenaar R, Young D, Cronin-Golomb A. Impact of optic flow perception and egocentric coordinates on veering in Parkinson’s disease. Brain 2008; 131: 2882–2893.

DeMaagd G, Philip A. Parkinson’s disease and its management part 1: Disease entity, risk factors, pathophysiology, clinical presentation, and diagnosis. P T 2015

Ernst MO, Banks MS. Humans integrate visual and haptic information in a statistically optimal fashion. Nature 2002; 415: 429–433.

Fernandez R, Duffy CJ. Early Alzheimer’s disease blocks responses to accelerating self-movement. Neurobiol. Aging 2012

Fernandez R, Monacelli A, Duffy CJ. Visual motion event related potentials distinguish aging and alzheimer’s disease. J. Alzheimer’s Dis. 2013

Fetsch CR, Turner AH, DeAngelis GC, Angelaki DE. Dynamic reweighting of visual and vestibular cues during self-motion perception. J. Neurosci 2009; 29: 15601–15612.

Fleming SM, Weil RS, Nagy Z, Dolan RJ, Rees G. Relating introspective accuracy to individual differences in brain structure. Science (80-.). 2010; 329: 1541–1543.

Funato T, Aoi S, Oshima H, Tsuchiya K. Variant and invariant patterns embedded in human locomotion through whole body kinematic coordination. Exp. Brain Res. 2010; 205: 497–511.

Gullett JM, Price CC, Nguyen P, Okun MS, Bauer RM, Bowers D. Reliability of three Benton Judgment of Line Orientation short forms in idiopathic Parkinson’s disease. Clin. Neuropsychol. 2013; 27: 1167–1178.

Halperin O, Israeli-Korn S, Yakubovich S, Hassin-Baer S, Zaidel A. Self-Motion Perception in Parkinson’s disease [Internet]. Eur. J. Neurosci. 2020[cited 2020 Mar 22] Available from: http://doi.wiley.com/10.1111/ejn.14716

Jacobs RA. Optimal integration of texture and motion cues to depth. Vision Res. 1999; 39: 3621–3629.

Jankovic J. Parkinson’s disease: clinical features and diagnosis. J. Neurol. Neurosurg. Psychiatry 2008; 79: 368–376.

Jankovic J. Movement disorders in 2016: Progress in Parkinson disease and other movement disorders. Nat. Rev. Neurol. 2017

Jaywant A, Shiffrar M, Roy S, Cronin-Golomb A. Impaired Perception of Biological Motion in Parkinson’s Disease. Neuropsychology 2016; 30: 720–730.

Kwon OS, Tadin D, Knill DC. Unifying account of visual motion and position perception. Proc. Natl. Acad. Sci. U. S. A. 2015; 112: 8142–8147.

Landy MS, Kojima H. Ideal cue combination for localizing texture-defined edges. J. Opt. Soc. Am. A Opt. Image Sci. Vis 2001; 18: 2307–2320.

Li Y, Guo S, Wang Y, Chen H. Altered motion repulsion in Alzheimer’s disease. Sci. Rep. 2017

Lin CC, Wagenaar RC, Young D, Saltzman EL, Ren X, Neargarder S, et al. Effects of Parkinson’s disease on optic flow perception for heading direction during navigation. Exp. brain Res. 2014; 232: 1343–1355.

Mak MKY, Pang MYC. Balance confidence and functional mobility are independently associated with falls in people with Parkinson’s disease. J. Neurol. 2009

Milne E, Swettenham J, Hansen P, Campbell R, Jeffries H, Plaisted K. High motion coherence thresholds in children with autism. J. Child Psychol. Psychiatry Allied Discip. 2002

Montse A, Pere V, Carme J, Francesc V, Eduardo T. Visuospatial deficits in parkinsons disease assessed by judgment of line orientation test: Error analyses and practice effects. J. Clin. Exp. Neuropsychol. 2001; 23: 592–598.

Morgan MJ, Ward R. Conditions for motion flow in dynamic visual noise. Vision Res. 1980

Nasreddine ZS, Phillips NA, BÃ©dirian V, Charbonneau S, Whitehead V, Collin I, et al. The Montreal Cognitive Assessment, MoCA: A Brief Screening Tool For Mild Cognitive Impairment. J. Am. Geriatr. Soc. 2005; 53: 695–699.

Peirce J, Gray JR, Simpson S, MacAskill M, Höchenberger R, Sogo H, et al. PsychoPy2: Experiments in behavior made easy. Behav. Res. Methods 2019; 51: 195–203.

Postuma RB, Berg D, Stern M, Poewe W, Olanow CWW, Oertel W, et al. MDS clinical diagnostic criteria for Parkinson’s disease. Mov. Disord. 2015; 30: 1591–1601.

Pouget A, Beck JM, Ma WJ, Latham PE. Probabilistic brains: knowns and unknowns. Nat. Neurosci. 2013; 16: 1170–1178.

Putcha D, Ross RS, Rosen ML, Norton DJ, Cronin-Golomb A, Somers DC, et al. Functional correlates of optic flow motion processing in Parkinson’s disease. Front. Integr. Neurosci. 2014; 8: 57.

Rajananda S, Lau H, Odegaard B. A random-dot kinematogram for web-based vision research. J. Open Res. Softw. 2018

Raposo D, Sheppard JPP, Schrater PRR, Churchland AKK. Multisensory decision-making in rats and humans. J. Neurosci 2012; 32: 3726–3735.

Robertson CE, Martin A, Baker CI, Baron-Cohen S. Atypical Integration of Motion Signals in Autism Spectrum Conditions. PLoS One 2012

Schütt HH, Harmeling S, Macke JH, Wichmann FA. Painfree and accurate Bayesian estimation of psychometric functions for (potentially) overdispersed data. Vision Res. 2016; 122: 105–123.

Shadlen MN, Newsome WT. Motion perception: Seeing and deciding. Proc. Natl. Acad. Sci. U. S. A. 1996; 93: 628–633.

Shulman LM, Taback RL, Rabinstein AA, Weiner WJ. Non-recognition of depression and other non-motor symptoms in Parkinson’s disease. Parkinsonism Relat. Disord. 2002; 8: 193–197.

Song Y, Wang H. Motion-induced position mis-localization predicts the severity of Alzheimer’s disease. J. Neuropsychol. 2019

Tomlinson CL, Stowe R, Patel S, Rick C, Gray R, Clarke CE. Systematic review of levodopa dose equivalency reporting in Parkinson’s disease. Mov. Disord. 2010

Vaugoyeau M, Viel S, Assaiante C, Amblard B, Azulay JP. Impaired vertical postural control and proprioceptive integration deficits in Parkinson’s disease. Neuroscience 2007; 146: 852–863.

Weil RS, Schrag AE, Warren JD, Crutch SJ, Lees AJ, Morris HR. Visual dysfunction in Parkinson’s disease. Brain 2016; 139: 2827–2843.

White TL, Sadikot AF, Djordjevic J. Metacognitive knowledge of olfactory dysfunction in Parkinson’s disease. Brain Cogn. 2016; 104: 1–6.

Williams DW, Sekuler R. Coherent global motion percepts from stochastic local motions. Vision Res. 1984; 24: 55–62.

Yakubovich S, Israeli-Korn S, Halperin O, Yahalom G, Hassin-Baer S, Zaidel A. Visual self-motion cues are impaired yet overweighted during visual–vestibular integration in Parkinson’s disease. Brain Commun. 2020; 2

Yuille AL, Grzywacz NM. A computational theory for the perception of coherent visual motion. Nature 1988; 333: 71–74.

Zaidel A, Turner AH, Angelaki DE. Multisensory calibration is independent of cue reliability. J. Neurosci. 2011; 31: 13949–13962.

